# Recombinant SARS-CoV-2 spike proteins for sero-surveillance and epitope mapping

**DOI:** 10.1101/2020.05.21.109298

**Authors:** Sophie M. Jegouic, Silvia Loureiro, Michelle Thom, Deepa Paliwal, Ian M. Jones

## Abstract

The newly emergent SARS-CoV-2 coronavirus is closely related to SARS-CoV which emerged in 2002. Studies on coronaviruses in general, and SARS in particular, have identified the virus spike protein (S) as being central to virus tropism, to the generation of a protective antibody response and to the unambiguous detection of past infections. As a result of this centrality SARS-CoV-2 S protein has a role in many aspects of research from vaccines to diagnostic tests. We describe a number of recombinant forms of SARS-CoV-2 S expressed in commonly available expression systems and their preliminary use in diagnostics and epitope mapping. These sources may find use in the current and future analysis of the virus and the Covid-19 disease it causes.

## Introduction

The appearance of Covid-19 in late 2019 and its subsequent development to a pandemic have been widely reported [1]. Bioinformatics shows that the causative virus, SARS-CoV-2, is closely related to SARS, a beta coronavirus that caused an epidemic in 2002/3 and probably emerged by zoonotic transfer from an animal species [2, 3]. The basis of immunity for many coronaviruses is the Spike protein (S), a 140kDa type I membrane glycoprotein found on the surface of the virus. The SARS and SARS-CoV-2 S proteins are >75% homologous at the amino acid level and have many features in common, not least the use of the same receptor, Angiotensin Converting Enzyme 2 (ACE2), for virus entry. Recently determined structures of S in complex with ACE2 have confirmed the same folded receptor binding domain (RBD) in both S proteins, albeit slightly offset when compared with each other [4, 5]. The SARS-CoV-2 S has a polybasic site upstream of the S fusion peptide and preliminary experiments show that proteolytic processing of S is required for cell entry [6, 7]. S is the principle neutralizing determinant of the virus and is composed of two domains, S1 and S2. The S1 domain encompasses the RBD while the S2 domain encodes the fusion peptide, heptad repeats and transmembrane domain. Many previous studies on SARS S have shown that while the full-length molecule is sufficient for protection [8], S1 or the RBD alone offer similar protection without the possibilities of enhancing antibodies which have been reported in some cases [9, 10]. Thus, S, or fragments of S encompassing the RBD, constitute candidate vaccines for SARS-Cov-2 as well as being useful for the detection and mapping of anti-S serum responses in convalescent or vaccinated individuals [11, 12]. We describe a range of SARS-CoV-2 S protein fragments produced in expression systems available in most laboratories which may find application for these purposes and for others investigating S structure-function relationships.

## Results

### Expression of SARS-CoV-2 S and S fragments

A full-length sequence verified SARS-CoV-2 S construct acted as template for a number of smaller fragments made by amplification of the requisite sequence by high fidelity PCR. These fragments were designed to encompass the known domains of S or to span the entire S coding region as ~200 amino acids overlapping by 100 amino acids (Figure 1).

**Figure 1.**
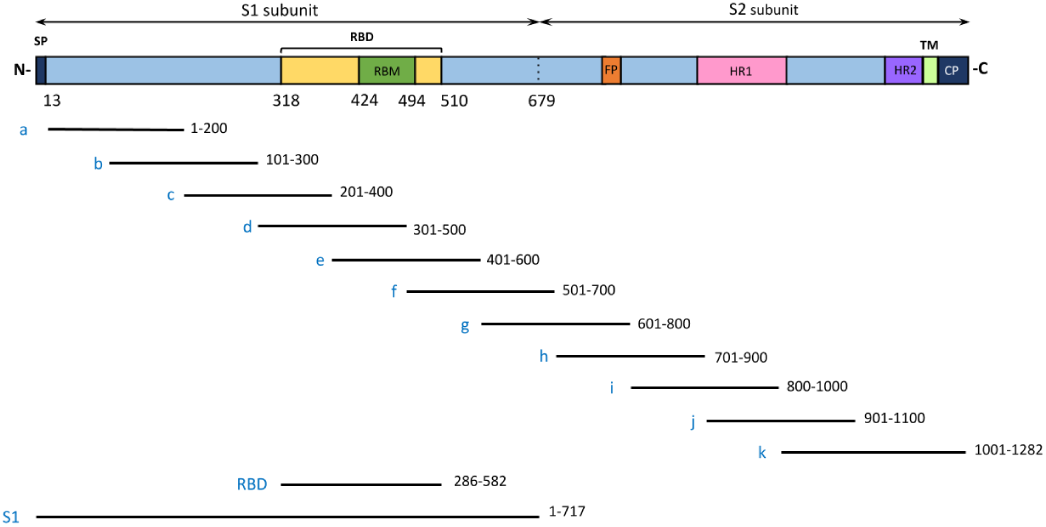
Cartoon representation of SARS-CoV-2 S protein with biological domains identified and the fragments used for S protein expression in insect cells and *E.coli*.

The original full length constructs and all smaller fragments were cloned into the multi-phylum expression vector pTriEx1.1 such that the encoded S sequence was appended at the C-terminus with a vector resident sequence encoding a polyhistidine tag for detection and purification, as used elsewhere [13]. To provide expression resilience and to facilitate different uses, two expression systems were used to generate recombinant S proteins and fragments thereof. Constructs encoding the full-length S and the S1 domain were transfected into Spodoptera frugiperda (Sf9) with linearized baculovirus DNA for the generation of recombinant baculoviruses [14]. In addition, all clones were transformed into BL21 related strains for T7 polymerase driven expression in *E.coli*.

Recombinant baculovirus stocks were used to infect small scale cultures (1 × 10^6^ cells) for confirmation of S protein expression. Analysis of infected insect cells and culture supernatant at 3 days post infection by western blot with an anti-His antibody showed expression of S and S1 in cell extracts and, as expected, S1 in the supernatant (Figure 2A). There was no evidence of an S2 band (~70kDa) in the S expressing cells suggesting that the S protein is not processed in insect cells, consistent with the requirement to engage the receptor [4] or incompatibility with insect cell furin [15]. Expression of S protein was also clear for full length S by immunofluorescence (permeabilized cells) or on the surface of cells as assessed by flow cytometry, both using an S reactive human monoclonal antibody (Figure 2 C & D). Larger scale cultures of S related proteins were expressed in *T.nao38* cells [16] and were purified following detergent lysis (S) or, in the case of S1, clarification of the infected cell supernatant. Single passage IMAC for the full length S protein enriched the non-glycosylated and glycosylated forms of the protein which were confirmed by western blot but also had a range of other insect cells proteins present (Figure 2B). However, direct IMAC of the S1 containing supernatant in the presence of 0.5mM nickel sulphate resulted in S1 that was ~85% pure (Figure 2B) with a yield of ~1mg/L (10^9^ cells).

**Figure 2.**
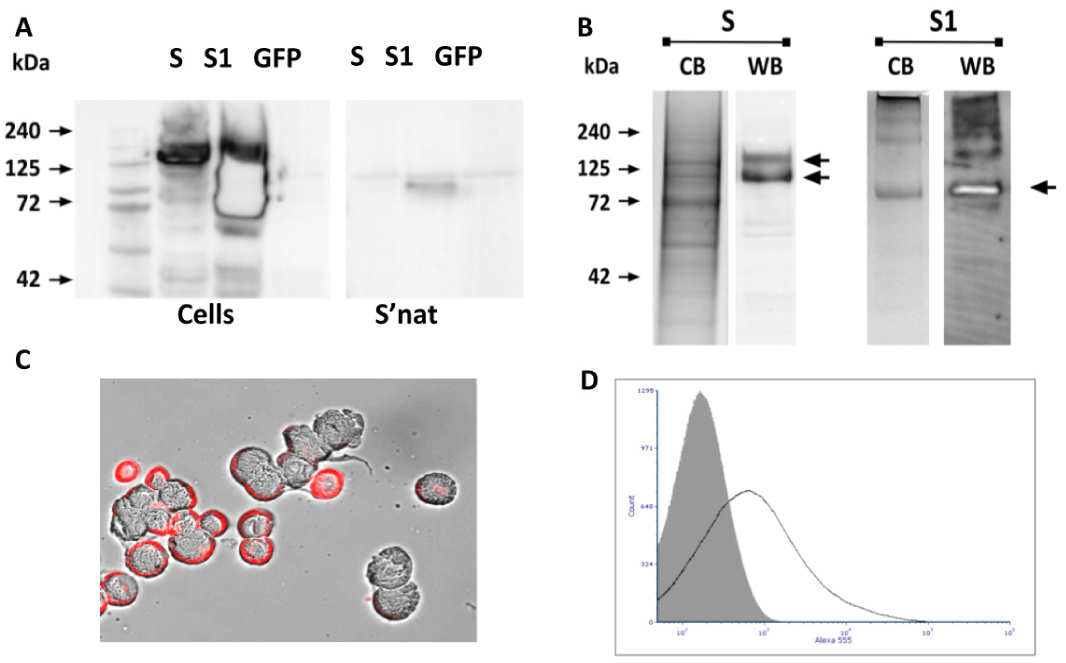
Expression of S and S1 in insect cells. A. Western blot of cells and supernatant with anti-His antibody. B. Purification of S and S1 by IMAC and confirmation of product by western blot. C. Immunofluorescence of S in permeabilized insect cells and D. flow cytometry of S expression on the surface of insect cells using CR3022 and an anti-human conjugate.

Similar western blot analysis of total protein extracts following induction of logarithmic phase *E.coli* cultures with IPTG confirmed the expression of His-tagged S antigen of the predicted molecular weight in all cases (Figure 3, upper panel). In general, the shorter overlapping set of S fragments, including those that spanned the RBD, were produced at higher levels than larger fragments following a 3 hr induction. Further analysis of the S-related proteins showed all well-expressed fragments to be produced as inclusion bodies which remained the case following low temperature induction (15°C). Retransformation of SoluBL21 (AMS Biotechnology) and LOBSTR [17] strains did not result in soluble protein expression despite titration of the IPTG concentration used for induction and further work is required on rescue strategies commonly in use to solubilize inclusion bodies (e.g. [18]). Nevertheless, we found that S protein fragments prepared for gel electrophoresis using non-reducing loading buffer could be used successfully for epitope mapping of 2 S reactive monoclonal antibodies, 3G9, an unpublished mouse mAb generated to SARS S, and CR3022, a human mAb also isolated originally to SARS [19] but shown to cross-react with SARS-CoV-2. A structure of the latter HuMAb in complex with the RBD has recently been solved [20]. By blot 3G9 bound to overlapping fragments c and d spanning resides 200-500, overlap 300-400 (Figure 3, middle panel) while CR3022 bound only to fragment c (Figure 3, lower panel). CR3022 was previously mapped to residues 369-519 which, in the modelled S trimer, are only exposed in certain conformations [20]. Our mapping suggests core binding by CR3022 can occur to a smaller region than that suggested by the structural footprint, within residues 369-400, in keeping with its reported non-sensitivity to mutation P462L [19].

**Figure 3.**
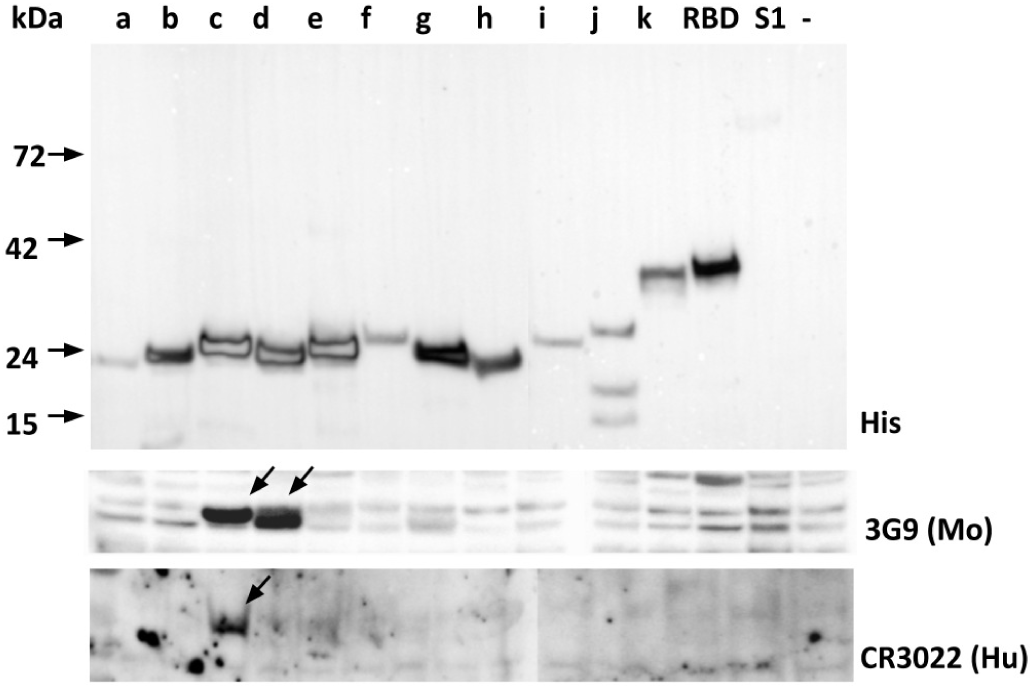
Expression of S protein fragments in *E.coli*. Induced cultures were resolved by SDS-PAGE and blotted with an anti His antibody (top panel), mouse monoclonal antibody 3G9 (middle panel) or CR3022 (lower panel). The lower two panels are cropped but showed no reaction to RBD or S1 despite their presence at detectable levels (upper panel).

### Configuration and use

The principal purpose in generating SARS-CoV-2 S protein or protein fragments were for studies of antibody binding or antibody generation. Accordingly, we used S1 expressed and purified from insect cells as an antigen for serum binding. 36 donated human sera from individuals, including some who had experienced Covid-19-like symptoms, were assessed for S reactivity. Preliminary titration experiments using an antibody to the His tag determined that S1 coating at 4μg per ml saturated the plastic surface. In addition, assays using CR3022 spiked into normal human plasma determined that, of several blocking buffers assessed, blocking the plate with steelhead salmon serum (Sea Block, Thermo Scientific), provided the lowest backgrounds. Standard ELISA of the sera with these conditions led to the identification of 5 sera (14%) as positive and 28 sera (77%) as negative and these were discriminated clearly at either the 1:40 or 1:80 dilution points (Figure 4, inset). Three sera gave intermediate binding curves and could not be unambiguously scored. The titration curves for the 5 positives were similar with anti S1 titres of ~500 in all cases (Figure 4).

**Figure 4.**
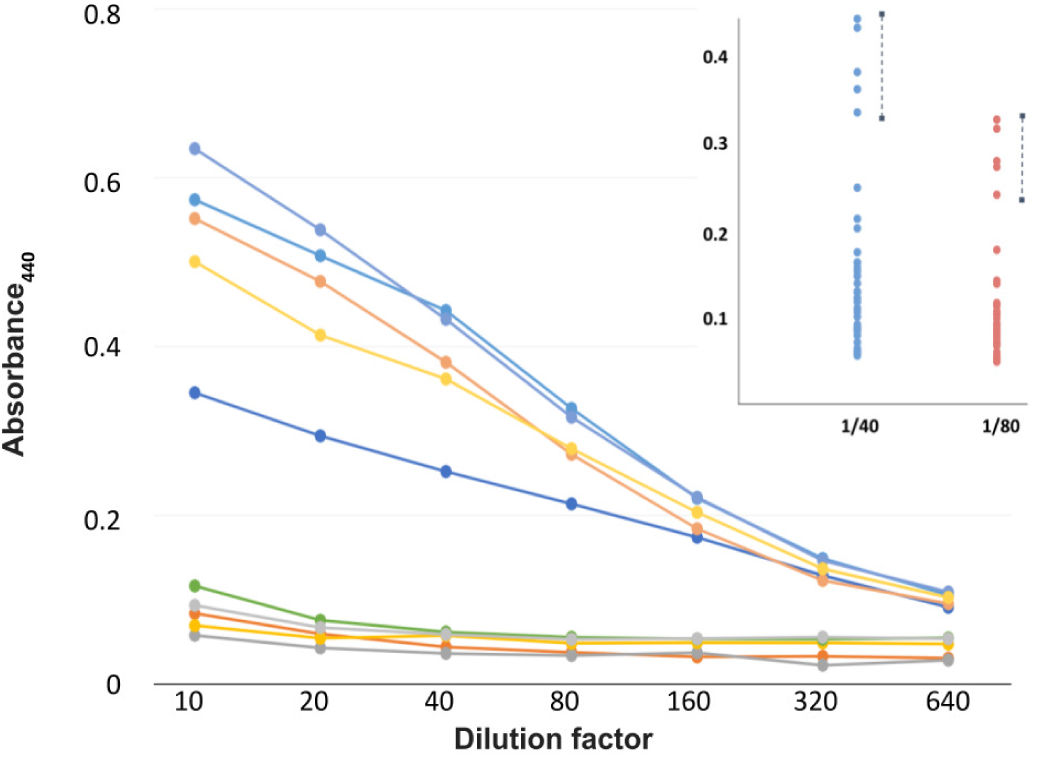
Screen for S1 reactivity. Sera were screened by ELISA on purified S1 protein starting at a dilution of 1:10. Inset: Scatterplot of all sera (n=36) at 1:40 and 1:80 dilution points with positives identified.

To provide an additional level of validation and to add epitope specificity to the data, 2 of the sera scoring positive by S1 ELISA were used as probes on western blots using full length S expressed in insect cells (*cf*. Figure 2) and also on the overlapping set of S fragments expressed in *E.coli* (*cf*. Figure 3). Both sera reacted with full length S antigen (Figure 5A) and showed binding to overlapping S fragments c and d, encompassing the RBD (Figure 5B). These data suggest that a second tier positivity test based on western blot could be used as confirmation of past infection and that, at least for antibodies able to bind to gel resolved antigen, antibodies to the RBD are present in convalescent individuals.

**Figure 5.**
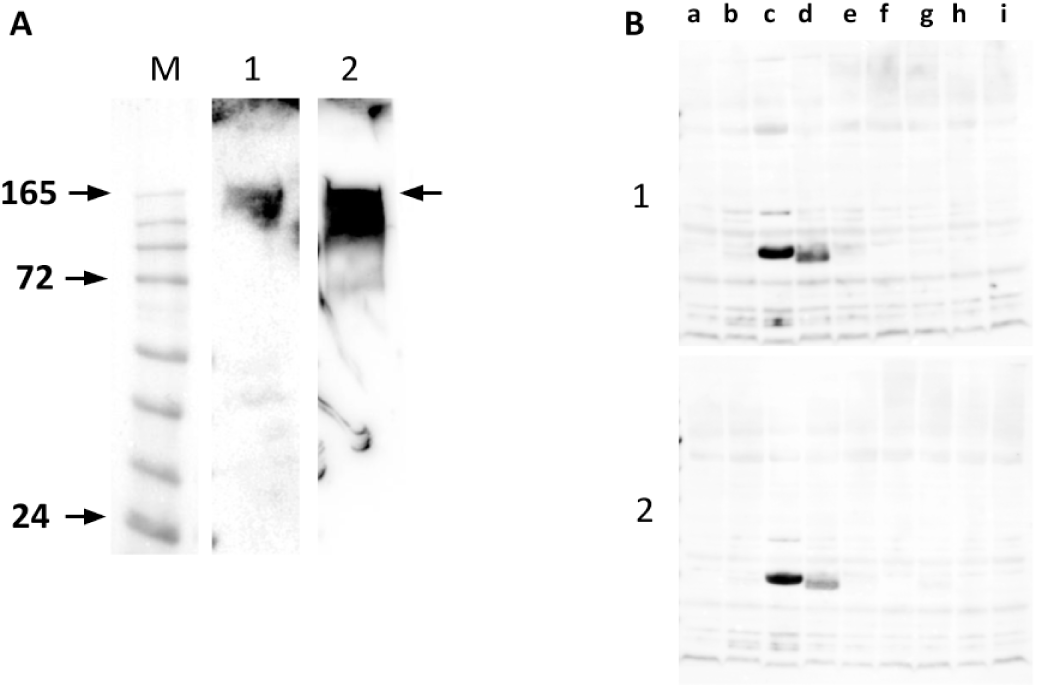
Western blot of two ELISA positive sera on full length S protein expressed in insect cells (A) or the overlapping set of S fragments as described in figure 3 (B).

## Discussion

The appearance of SARS-CoV-2 and its pandemic spread has led to the reported expression of the virus encoded proteins, notably the S protein, for structural study [4, 5], for immunisation [21] and for diagnostics [22]. We have described recombinant sources of several S related polypeptides from two common expression systems, recombinant baculoviruses and *E.coli* and used the proteins expressed for the analysis of seroconversion and for epitope mapping. The particular properties of the insect cell system, yield, robustness and the ability to perform at scale are discussed elsewhere [23], similarly the use of the T7 system in *E.coli [24]*. Using a set of overlapping S fragments we demonstrated epitope mapping of monoclonal antibodies 3G9 and CR3022 to residues 300-400 of S. Interestingly, neither antibody reacted with the RBD itself in the western blot format used despite it encompassing the residues recognised. This suggests that each fragment of S may adopt a variable level of refolding on the membrane following transfer and emphasizes the value of the overlapping fragment approach to enable epitope identification. Of 28 residues in the RBD shown to interact with CR3022, 20 lie within residues 369-393 [20] and this core region alone is evidently sufficient to allow binding, as shown by interaction with S fragment c here. The overlapping fragment set allowed mapping of two human sera which also showed binding to the same region. It remains to be determined how widespread this serum response is in exposed individuals and if there is any correlation between the epitope specificity of a serum and the titre or level of neutralization.

The globular S1 domain of several coronaviruses is the preferred antigen for sero-surveillance [6, 22, 25, 26] and purified SARS-CoV-2 S1 was used similarly here. Full length S protein has been used elsewhere [11] but the S2 domain which it includes can lead to cross reaction as a result of previous coronavirus infections [27]. Although we used purified S1, we have shown previously that glycosylated antigen from infected insect cells captured to the plate by a mannose specific lectin (GNA) can also be an effective antigen, avoiding the need for protein purification in resource limited situations [28]. We found evidence for ~14% seroconversion in a set of random samples, some of which were confirmed by western blot. The titre of all these sera was similar and as no other information on the samples was available no correlation with symptoms was possible although none of the samples were from hospitalised individuals. The level of seroconversion in the UK population is currently unknown although unpublished data suggest a range of 5-10% [29]. Oxfordshire has a demonstrated positivity rate of 269 per 100,000 (Office for National Statistics) which, using a 50-fold factor for non-tested but exposed individuals suggested from other studies [30], would suggest an infection rate of ~13%, very close to our observed positivity. Several studies of seroconversion have been reported, in hospitalised individuals [22] and in the population generally [27] but the general relationships among disease severity, antibody titre, neutralisation, epitope profile and longevity of response remain to be determined. The set of S resources described here may contribute to studies in these areas.

## Materials and Methods

### Constructs

The sequence of SARS-CoV-2 S (accession no. NC_045512) was codon optimised for *Spodoptera frugiperda* cells and the resident signal peptide exchanged for that of honeybee melittin [31] before being ordered as two overlapping fragments (IDT Europe) flanked by 18bps at the 5’ and 3’ ends homologous to the intended expression vector, pTriEx1.1 (EMD Millipore). The 3’ flanking nucleotides were also designed to fuse the S open reading frame in-frame to the vector resident polyhistidine encoding sequence (6 His residues). The gene was assembled at the same time as recombination into the vector using infusion technology (NEB) and the assembly reaction used to transform *E.coli* (NEB 10 beta). Colonies positive by PCR screening using primers that flank the cloning site were sequenced across the entire S coding region and a single positive isolate adopted for all further manipulations.

### Cell culture

*Spodoptera frugiperda* (Sf9) and *T.niao*38 cells were maintained in EX-CELL 420 medium (Sigma) supplemented with 2% fetal bovine serum, 1% penicillin/streptomycin solution, at 27°C with shaking. Virus growth used exclusively *Sf*9s while protein expression used predominantly *T.niao38*.

### Baculovirus expression

Linearized baculovirus DNA was used to produce recombinant baculoviruses [14]. Small-scale protein expression was performed by infection of a 6-well plate seeded with 1 × 10^6^ Sf9 cells per well using 200μl of a high titre stock of the recombinant baculovirus, typically passage 3, and incubated for 5 days at 27°C. After incubation, cells were harvested and used for antigen detection.

### E.coli expression

Constructs were transformed into *E.coli* T7 Express lysY (NEB) and isolated by ampicillin resistance. Cultures were grown at 37°C to an OD600=0.6 and induced by the addition of IPTG to 0.4mM. Growth was continued at 27°C for 3 hr and cells harvested and disrupted for gel or purification as required. SDS-PAGE analysis used the equivalent of 50 μl of culture per lane. Solubility was gauged after lysis in Bugbuster (EMD) and gel analysis of the soluble and pellet fractions. Alternate hosts used were SoluBL21 (AMS Biotechnology) and LOBSTR [17].

### Immunofluorescence

Forty-eight hours after infection, cells were dislodged by pipette and washed twice with cold PBS for 5 minutes, then incubated in fix and permeabilization buffers for 1hr at room temperature (eBioscience™). Fixed and permeabilized cells were incubated with CR3022 at 2 μg per ml for 1 hr at room temperature. They were washed in PBS 3% BSA and then incubated with anti-human AlexaFluor 555 conjugate for a further 1hr. The cells were washed twice with TBS for 15 min each at room temperature mounted with a drop of Slowfade™ Gold reagent before being imaged using an EVOS-FL digital fluorescence microscope (Thermo Fisher Scientific). Cells for flow cytometry were processed similarly, but without permeabilization, and data acquired using a BD Accuri C6 Plus and analysed by FC Express v7 (De Novo software).

### Protein Purification

Infected insect cells were disrupted with CytoBuster™ protein extraction reagent (Merck) and clarified by centrifugation at 4,300 × g for 20m before column loading. For S1, the supernatant of infected cells was clarified by centrifugation as above followed by passage through a 0.8 micro filter. The clear supernatant was adjusted to 0.5mM nickel sulphate to avoid stripping the IMAC column before loading. In both cases IMAC chromatography was done using a pre-prepared 5ml IMAC column (GE) operating at a flow rate of 2.5 ml.min^−1^ with a gradient elution of 0.05-0.25M imidazole over 60 minutes.

### SDS-PAGE

Proteins were disrupted in NuPAGE loading buffer (Thermo) and separated by SDS-PAGE using 4-12% precast Tris-Glycine SDS polyacrylamide gels (Invitrogen) for 30min at 200V. After electrophoresis, gels were either stained with Coomassie blue R250 or transferred to PVDF membranes for Western blot analysis. For epitope mapping using overlapping S fragments, inclusion bodies were disrupted in 3% SDS loading buffer without reducing agent.

### Western blot

Following semi-dry transfer to PVDF membranes, membranes were incubated in TBST blocking buffer consisting of (5% of skimmed milk powder, 0.2% Tween-20, 1 × TBS) for 1 hr. Membranes were incubated with the primary antibody at 1:1,000 in 1x TBST buffer for 1hr followed by 3 × washes for 5 minutes each in TBST buffer and if necessary with a secondary horseradish-peroxidase (HRP) antibody conjugate (Sigma) diluted 1:1,000 in 1x TBST for 1hr followed by 3 × washes for 5 minutes each with TBST buffer. The membrane was finally washed with TBS and HRP activity detected using chemiluminescence imagery.

### ELISA

Microtitre plates (Nunc Maxisorb) were coated with S1 antigen at 4μg per ml in 50mM sodium carbonate-bicarbonate buffer (pH 9.6) for a minimum of 1 hour at room temperature. Excess antigen was removed by washing three times with Super block (Thermo Scientific) and unbound sites blocked using non-diluted Sea Block. Samples were added at 1/10 dilution and diluted in a 2-fold series thereafter, followed by 1 hr at room temperature. The plates were washed five times with TBS containing 0.05% v/v Tween-20 and polyclonal anti-human antibody conjugated to HRP (Sigma) added for one hour at room temperature. Following washing the chromogenic substrate TMB (Europa Bioproducts) was added and colour development stopped by the addition of a 40% well volume of 0.25M sulphuric acid. Absorbance was read at 440nm against a reference read at 700nm.

### Blood samples

Finger prick samples from volunteers contacted by word of mouth were collected into 4% sodium citrate using self-retracting lancets. The samples were collected in the last week of April 2020 in central Oxfordshire. No other information on the donated sample was sought.

## End Matter

### Author Contributions and Notes

IMJ designed research, SMJ, SL, MT, DP and IMJ performed research and analysed data; and IMJ and SMJ wrote the paper.

The authors declare no conflict of interest.

Reagent requests should be made to IMJ.

## Acknowledgments

We acknowledge the Williams family and their anonymous donors for their help in enabling the serology. Phil Lowry kindly read the manuscript.

